# Orai1-mediated Ca^2+^ Entry Regulates Lipolysis and Mitochondrial Activation in Brown Adipose Thermogenesis

**DOI:** 10.64898/2026.04.24.718619

**Authors:** Suji Kim, Nguyen Phan Anh, Kyu-sang Park, Seung-Kuy Cha, Jun Namkung

## Abstract

Cold-induced thermogenesis in brown adipose tissue (BAT) is essential for maintaining energy homeostasis, yet the Ca^2+^-dependent mechanisms underlying this process remain incompletely understood. Here, we identify Orai1, a component of the store-operated Ca^2+^ entry pathway, as a regulator of thermogenic activation in BAT. Using a brown adipocyte-specific *Orai1* knockout mouse model, we demonstrate that cold exposure is associated with Orai1-dependent Ca^2+^ influx through a non-canonical mechanism. Orai1 deficiency leads to impaired cAMP-PKA signaling, reduces the expression of lipolytic enzymes and thermogenic genes, and diminished mitochondrial Ca^2+^ uptake and uncoupling. These defects culminate in cold intolerance, lipid accumulation, and decreased energy expenditure. Mechanistically, Orai1 facilitates Ca^2+^-dependent activation of adenylyl cyclase 3, linking membrane Ca^2+^ entry to cAMP production, and promotes mitochondrial remodeling and oxidative metabolism. These findings support a key role for Orai1 in coordinating Ca^2+^ entry to lipolytic and mitochondrial pathways in brown adipocytes and highlight its potential therapeutic target in metabolic diseases characterized by impaired energy metabolism.

**HIGHLIGHTS:**

- Orai1 mediates Ca^2+^ influx in brown adipocytes through a non-canonical, partially STIM1-independent mechanism.
- Orai1-mediated Ca^2+^ influx promotes both cAMP-PKA-driven lipolysis and mitochondrial oxidative activation.
- Orai1-dependent Ca^2+^ entry promotes cAMP-PKA signaling and lipolytic activation I nbrown adipocytes.
- Orai1 coordinates mitochondrial Ca^2+^ uptake to support thermogenic function in brown adipocytes.

## INTRODUCTION

Obesity and its associated metabolic disorders – such as type 2 diabetes, dyslipidemia, and fatty liver disease – remain major public health burdens despite advances in pharmacological and lifestyle interventions^1^. Among the various strategies to counteract excess energy store, enhancing energy expenditure via brown adipose tissue (BAT) has gained renewed attention. In adult humans, BAT activity inversely correlates with adiposity and insulin resistance, and its stimulation improves systemic metabolic profiles, including glucose and lipid homeostasis^2,3^.

BAT exerts a pivotal role in non-shivering thermogenesis, which is induced by cold exposure to generate heat through mitochondrial uncoupling mediated by uncoupling protein 1 (UCP1)^4-6^. This thermogenic response is classically attributed to sympathetic activation and β-adrenergic signaling, which promote lipolysis and mitochondrial respiration via cAMP-protein kinase A (PKA) pathway^7^. In parallel, thermosensitive Ca^2+^-permeable channels such as TRPV1, TRPM8, and TRPA1, predominantly studied in sensory neurons, have been implicated in temperature perception and energy homeostasis^8,9^

While the central role of cAMP signaling in cold-induced BAT activation is well established, emerging evidence suggests that Ca^2+^ signaling also serves as an essential integrator of metabolic activation in brown adipocytes. Specifically, mitochondrial Ca^2+^ uptake boosts oxidative metabolism and thermogenic gene expression, whereas cytosolic Ca^2+^ can stimulate adenylyl cyclases (ACs) to amplify lipolytic signaling^10,11^. However, the molecular identity of Ca^2+^ influx pathways during thermogenic activation remains elusive, and the extent to which Ca^2+^-dependent signaling to thermogenic regulation is poorly understood.

The Orai1-STIM1 complex, canonically known for mediating store-operated Ca^2+^ entry (SOCE)^12-15^, has recently been implicated in temperature-dependent Ca^2+^ signaling. STIM1 undergoes temperature-dependent conformational changes, functioning as a heat sensor in keratinocytes, while the Orai1-STIM1 complex has been shown to be associated with cold-triggered Ca^2+^ influx in sensory and sympathetic neurons^16-18^. However, whether Orai1 plays a functional role in thermogenic regulation in brown adipocyte remains unknown.

In this study, we used a brown adipocyte-specific *Orai1* knockout (*Orai1* BKO) model and complementary *in vitro* approaches to investigate the role of Orai1 in brown adipocytes. We show that Orai1-mediated Ca^2+^ influx promotes cAMP production and lipolysis via AC activation, and mitochondrial Ca^2+^ uptake for thermogenesis. Loss of Orai1 impairs these processes and reduces thermogenic capacity in brown adipose tissue. Our findings support a role for Orai1 in coordinationg Ca^2+^-dependent signaling pathways that enable thermogenic activation.

## MATERIALS AND METHODS

### Animals

All mouse experiments were conducted under protocols approved by the Institutional Animal Care and Use Committee at Yonsei University Wonju College of Medicine (Approval number: YWC-181022-1). Brown adipocyte-specific *Orai1* knockout mice (*Orai1*^*fl/fl*^, *Ucp1-Cre*^*+*^) were generated by crossing *Orai1* floxed mice^19^ (kindly provided from Dr. Yousang Gwack, UCLA, USA) with *Ucp1* promoter-driven *Cre* transgenic mice (the Jackson Laboratory 024670). Littermates without Cre (*Orai1*^*fl/fl*^, *Ucp1-Cre*^*-*^) were used as control. Male littermates (8-12 weeks) were used for all experiments. Mice were housed at 22 ± 1 °C under a 12-hour light/dark cycle with *ad libitum* access to food and water. For high-fat diet (HFD) studies, mice were fed a 60 % kcal fat diet (D12492, Research Diets) for 12 weeks. Body mass was measured using nuclear magnetic resonance (LF50, Bruker). Cold exposure was performed by placing mice in a 4 °C chamber for the indicated durations. Core body temperature was monitored using implantable telemetry probes (G2 E-Mitter, STARR LifeScience), and skin temperature was measured using a thermal imaging camera (FLIR T530). Indirect calorimetry (O_2_ consumption, CO_2_ production, RER, and heat production) was assessed using a Comprehensive Lab Animal Monitoring System (Columbus Instruments). Glucose and insulin tolerance tests were performed following 6-hour fast. Blood glucose levels were measured from tail vein blood using a glucometer. Serum triglycerides (TG), total cholesterol (TC), HDL, and LDL levels were measured using enzymatic assays (Abnova).

### Cell cultures

Primary brown preadipocytes were isolated from interscapular BAT of control and *Orai1* BKO mice. Cells were cultured in DMEM/F12 supplemented with 10 % FBS and induced to differentiate using an adipogenic cocktail (0.5 mM IBMX, 1 μM dexamethasone, 5 μg/mL insulin, 1 nM T3, 125 μM indomethacin). Fully differentiated adipocytes (day 8) were used for *in vitro* experiments. hTERT-immortalized human brown preadipocytes (ATCC-CRL-4062) were maintained in DMEM/F12 medium supplemented with 10 % FBS. For differentiation, confluent cells were induced using a brown adipogenic differentiation cocktail as described above. All experiments using human brown adipocytes were performed after day 12 of differentiation. For knockdown experiments, mouse primary brown adipocytes were transfected after 5 days of differentiation, whereas human brown adipocytes were transfected after 12 days of differentiation. Cells were transfected with siRNAs targeting *Orai1, Stim1, Trpv2, and Trpm8* (Bioneer) using Lipofectamine RNAiMAX (Thermo Fisher). Cells were treated with forskolin (10 μM, Sigma), CL-316,243 (1 μM, Sigma), or norepinephrine (0.1 μM, Sigma) as indicated. Recombinant lentiviruses were generated by co-transfecting pLenti-CMV-Orai1-PuroR with pMDG.2 and psPAX2 into HEK293FT cells using X-tremeGENE™ HP DNA Transfection Reagent (Roche). Viral supernatants were collected 72 hours after transfection, filtered, concentrated using Lenti-X concentrator (TaKaRa), and stored at −80 °C. Differentiated cells were infected with lentivirus in complete culture medium, and the medium was replaced with fresh medium after 24 h.

### Intracellular Ca^2+^ imaging

Intracellular Ca^2+^ ([Ca^2+^]_i_) measurement using Fura-2 AM (Invitrogen) was performed as described previously^20,21^. The normal physiological salt solution (NPSS) used for bath perfusion contained 135 mM NaCl, 5 mM KCl, 1mM MgCl_2_, 2 mM CaCl_2_, 10 mM HEPES, and 5.5 mM glucose (pH 7.4). For SOCE measurements, cells were pretreated with cyclopiazonic acid (CPA, 20 μM) in Ca^2+^-free PSS followed by addition of NPSS. Fluorescence emission ratios (340/380 nm excitation) were recorded using a fluorescence imaging system (Olympus) with MetaFluor 6.1 software (Sutter Instrument). All [Ca^2+^]_i_ measurements were performed at 37 °C using a heat controller (Warner Instruments). Cold-induced [Ca^2+^]_i_ was assessed by rapidly changing bath perfusion media temperature from 37 °C to 15 °C.

### STIM1-ORAI1 puncta formation assay

T-REx 293 cells stably expressing STIM1-mCherry-T2A-ORAI1-GFP^22^ were used to visualize STIM1-ORAI1 puncta formation. Cells were incubated in NPSS containing 2 mM Ca^2+^ for 10 min at room temperature. To induce store-operated calcium entry as a positive control, cells were treated with CPA in Ca^2+^-free PSS for 10 min. For cold stimulation, cells were incubated in NPSS containing 2 mM Ca^2+^ at 15 °C for 10 min. Cells were fixed with 4% PFA, stained with DAPI, and analyzed for STIM1-ORAI1 puncta formation by fluorescence microscopy.

### Mitochondrial Ca^2+^ Uptake

Mitochondrial Ca^2+^ was assessed using Rhod-2 AM (Thermo Fisher), a mitochondria-targeted Ca^2+^ indicator. Fluorescence signals were recorded using a temperature-controlled imaging system (Flexstation 3) under temperature-controlled perfusion.

### Western Blotting

Protein lysates were extracted from BAT or cultured cells using RIPA buffer containing protease and phosphatase inhibitors. Equal amounts of protein were separated by SDS-PAGE and transferred to PVDF membranes. Antibodies used in this study are listed in Supplementary Table 1. Detection was performed using enhanced chemiluminescence (ECL, Thermo).

### qRT-PCR

Total RNA was extracted using Trizol reagent (Invitrogen). cDNA synthesis was performed with SuperScript III (Invitrogen). qRT-PCR was carried out using SYBR Green Master Mix (Applied Biosystems) on a QuantStudio 6 Real-Time PCR System (Thermo Fisher). Relative expression was calculated using ddCt method and normalized to *Rplp0*. Primer sequences are provided in Supplementary Table 2.

### Histology and Electron Microscopy

BAT, eWAT, iWAT, and liver tissues were fixed in 10% formalin, paraffin-embedded, sectioned, and stained with hematoxylin and eosin (H&E). For electron microscopy, BAT was fixed in 2.5% glutaraldehyde, post-fixed in osmium tetroxide, dehydrated, embedded, and sectioned. Images were acquired using transmission electron microscope (Hitachi).

### Mitochondrial function and DNA content

Primary brown adipocytes were assessed for metabolic function using a Seahorse XF Analyzer (Agilent). A mitochondrial stress test was performed with sequential injection of oligomycin, FCCP (carbonyl cyanide-4-(trifluoromethoxy)phenylhydrazone)), and a mixture of rotenone and antimycin A. Fatty acid oxidation (FAO) capacity was evaluated with or without prior treatment with etomoxir (40uM). Mitochondrial DNA copy number was assessed by qPCR for *mt-Nd1* and normalized to *Gapdh*.

### cAMP assay

Intracellular cAMP concentrations were quantified using a cAMP ELISA kit (Abcam, 138880) according to the manufacturer’s instructions. BAT tissues were homogenized in 0.1 M HCl to prevent phosphodiesterase activity before analysis.

### Statistical analysis

All data are expressed as mean ± standard error of the mean (SEM). Statistical analyses were conducted using GraphPad Prism (version 10). Differences between two groups were assessed using an unpaired, two-tailed Student’s t-test, while comparisons among multiple groups were analyzed by one-way analysis of variance (ANOVA). Statistical significance was indicated as follows: *p < 0.05; **p < 0.01; ***p < 0.001; ****p < 0.0001.

## RESULTS

### Orai1 mediates Ca^2+^ entry in brown adipocytes under thermogenic condition

BAT plays a central role in non-shivering thermogenesis, which is rapidly activated upon cold exposure^23^. To assess whether cold temperature elevates [Ca^2+^]_i_ via Orai1 (Fig. 1a), we performed live-cell Ca^2+^ imaging using a controlled temperature ramp (37 °C to 15 °C) in primary brown adipocytes. Acute cold exposure elicited a robust rise in [Ca^2+^]_i_ levels (Fig. 1b), suggesting that cold exposure is associated with increased Ca^2+^ influx in brown adipocytes. To directly compare Orai1 with other candidate thermosensitive Ca^2+^ channels, including Trpv2 and Trpm8^24,25^, we quantified cytosolic Ca^2+^ entry after siRNA-mediated knockdown in primary brown adipocytes (Supplementary Fig. 1a, b). In contrast to siOrai1, siTrpv2 and siTrpm8 did not suppress cold-induced Ca^2+^ responses relative to control cells. The Orai1-STIM1 complex, a key component of SOCE, has recently been shown to mediate cold-triggered Ca^2+^ influx in sensory and sympathetic neurons^16^. In a reconstituted STIM1-Orai1 imaging assay (Fig. 1c), CPA promoted canonical co-puncta and oligomerization, whereas acute cold increased STIM1 and Orai1 puncta without enhancing STIM1-Orai1 oligomerization (Fig. 1d), suggesting a non-canonical mode of Orai1 activation under these conditions. To investigate the role of Orai1 in Ca^2+^ entry under thermogenic stimulation in brown adipocytes, we generated a brown adipocyte-specific *Orai1* knockout (*Orai1* BKO; *Orai1*^*fl/fl*^, *Ucp1-Cre*^*+*^) (Fig. 1e). Primary brown adipocytes isolated from *Orai1* BKO exhibited markedly reduced SOCE in response to CPA-induced store depletion (Fig. 1f), validating Orai1 as the principal Ca^2+^ release-activated Ca^2+^ (CRAC) channel in these cells.

**Fig. 1.**
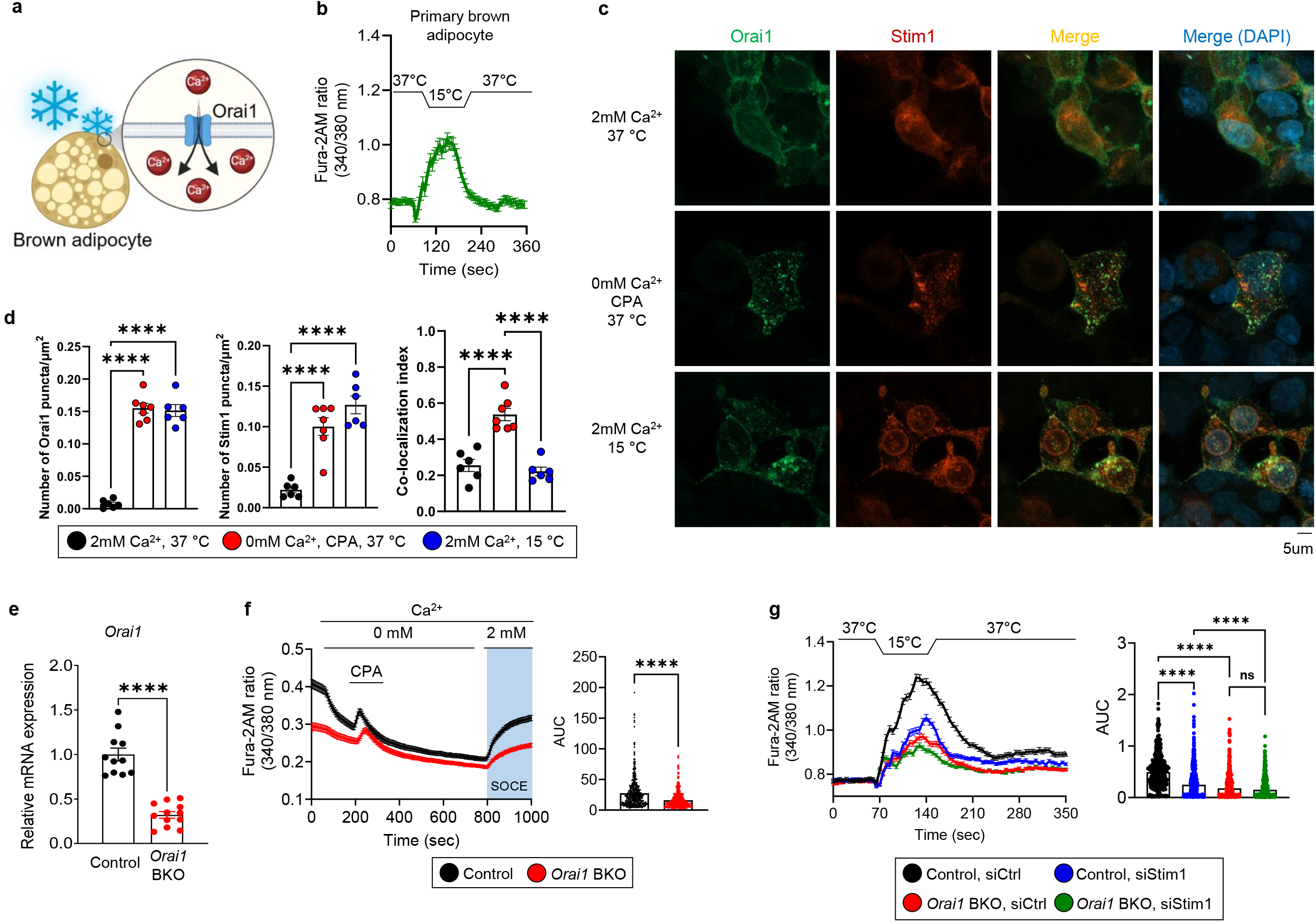
Orai1 contributes to cold-associated Ca^2+^ influx in brown adipocytes. **a** Schematic of the proposed model in which cold exposure is associated with Orai1-mediated Ca^2+^ influx, triggering thermogenic activation in brown adipocytes. **b** Intracellular Ca^2+^ levels measured by Fura-2 AM fluorescence in primary brown adipocytes following temperature shift from 37 °C to 15 °C. **c** Representative confocal images of HEK293T cells co-expressing STIM1-mCherry and Orai1-GFP at 37 °C, 15 °C, and CPA. **d** Quantification of puncta density and oligomerization. **e** Confirmation of *Orai1* deletion in BAT by qRT-PCR. **f** Store-operated Ca^2+^ entry in primary brown adipocytes from control and *Orai1* BKO mice measured by CPA-induced Ca^2+^ depletion/readdition assay. **g** Cold-induced Ca^2+^ influx in control and *Orai1* BKO adipocytes with or without *Stim1* knockdown. Data are presented as mean ± SEM. Statistical significance: \*\*\*\**p* < 0.0001.

We next examined cold-induced Ca^2+^ responses in control and *Orai1* BKO brown adipocytes. Acute cold exposure triggered significant Ca^2+^ influx in control cells, which was blunted in Orai1-deficient brown adipocytes (Fig. 1g). Knockdown of *Stim1* similarly reduced cold-induced Ca^2+^ influx in control cells, but had no further effect in *Orai1* BKO cells. These results suggest that Orai1 contributes to Ca^2+^ entry under these conditions, with a limited contribution from STIM1.

To determine whether cold exposure transcriptionally modulates *Orai1* expression, we assessed Orai1 mRNA levels in BAT under acute (6 hours) or chronic (3 days) cold exposure. Both conditions resulted in a significant upregulation of *Orai1* expression (Supplementary Fig. 1c, d), suggesting that Orai1 expression is regulated during thermogenic adapation. These findings indicates that Orai1 contributes to Ca^2+^ entry in brown adipocytes and is transcriptionally upregulated during thermogenic adaptation, supporting a role for Orai1 in Ca^2+^-dependent intracellular signaling under these conditions.

### *Orai1* deletion impairs energy expenditure and aggravates metabolic dysfunction

Given that Orai1 mediates Ca^2+^ influx in brown adipocytes, we next assessed whether its loss affects systemic energy metabolism in vivo. Under standard chow conditions, *Orai1* BKO mice exhibited increased fat mass compared to littermate control, without significant changes in body weight (Fig. 2a, b). Indirect calorimetry revealed a marked decrease in oxygen consumption and heat production, with no change in locomotor activity (Fig. 2c), suggesting that energy expenditure, rather than behavior, was impaired. Notably, these alterations were observed under baseline conditions, indicating reduced thermogenic capacity independent of acute stimulation. To assess whether *Orai1* deficiency increases susceptibility to metabolic stress, mice were subjected to high-fat diet (HFD) feeding. *Orai1* BKO mice exhibited accelerated weight gain compared to controls (Fig. 2d), along with exacerbated glucose intolerance and insulin resistance (Fig. 2e, f). Histological analysis revealed increased lipid accumulation in BAT and marked hepatic steatosis (Fig. 2g), while morphology of iWAT and eWAT remained largely unchanged (Supplementary Fig. 2). Consistent with metabolic dysfunction, serum levels of triglycerides and cholesterol were significantly elevated in *Orai1* BKO mice under HFD (Fig. 2h). Furthermore, expression of thermogenic genes such as *Ucp1, Dio2, Ppargc1a*, and *Prdm16* was significantly downregulated in BAT from *Orai1* BKO under HFD conditions (Fig. 2i), indicating impaired transcriptional activation of the thermogenic program. Together, these findings suggest that loss of Orai1 is associated with reduced basal thermogenic capacity and impaired energy expenditure, leading to increased vulnerability to obesity and metabolic dysfunction under nutritional overload.

**Fig. 2.**
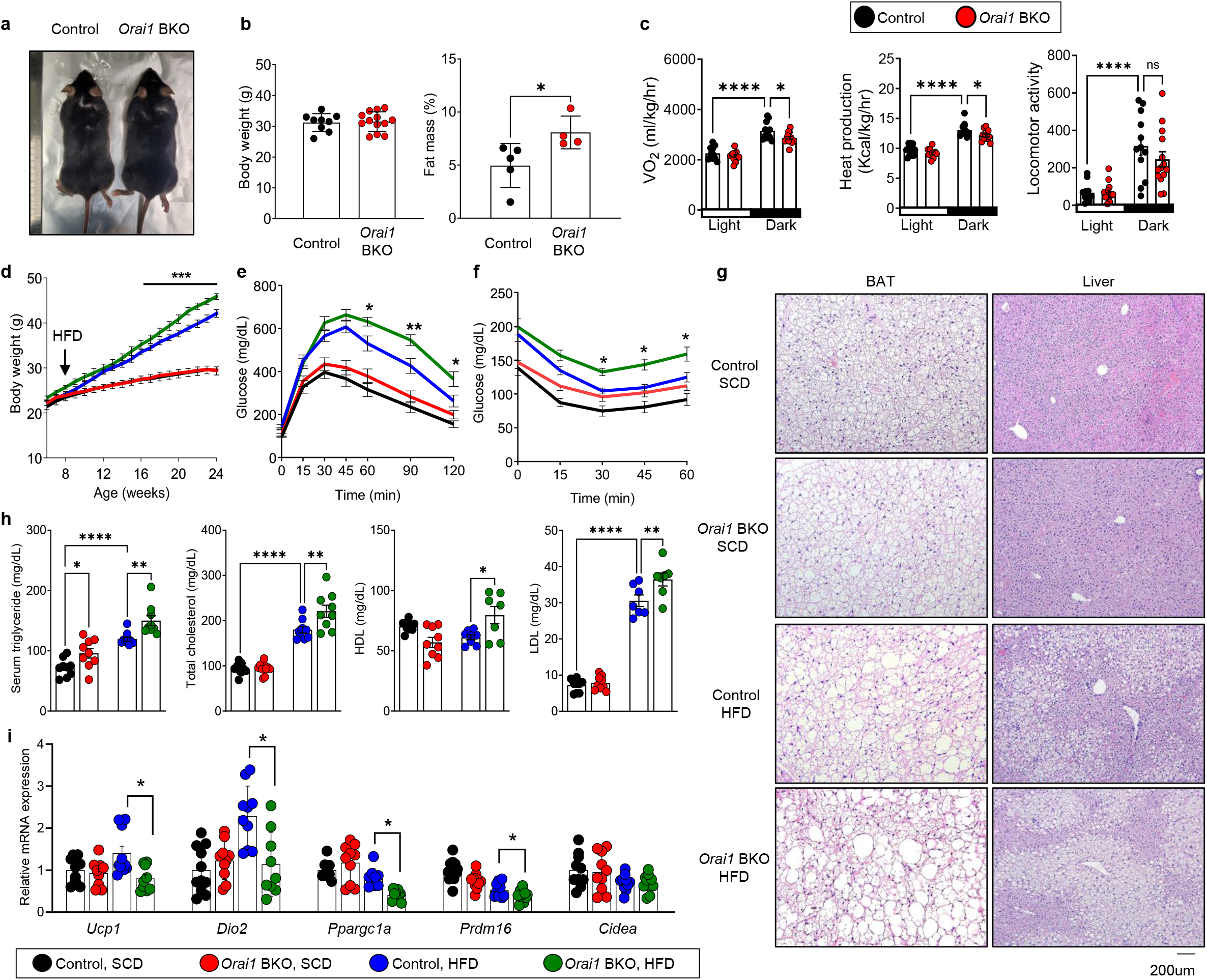
Loss of Orai1 in brown adipocytes leads to adiposity and metabolic dysfunction under high-fat diet (HFD). **a** Gross morphology of control and *Orai1* BKO mice (30 weeks). **b** Body composition analysis by NMR of control and *Orai1* BKO mice (30 weeks). **c** indirect calorimetry measurements (20 weeks). **d** Body weight curves under HFD conditions (n=11~13 mice per group). **e, f** Glucose tolerance test (**e**) and insulin tolerance test (**f**) in control and *Orai1* BKO mice fed HFD. **g** Representative H&E staining images of BAT and liver. **h** Serum biochemical analysis of lipid profiles. **i** qRT-PCR analysis of thermogenic genes in BAT. Data are presented as mean ± SEM. Statistical significance: \**p* < 0.05, \*\**p* < 0.01, \*\*\**p* < 0.001, \*\*\*\**p* < 0.0001.

### Orai1 deficiency impairs thermogenic responses and lipid mobilization under cold exposure

Given that *Orai1*-deficient mice exhibit reduced basal thermogenic activity, we next examined whether their impaired response to cold exposure reflects a primary sensing defect or a limitation in thermogenic capacity. We subjected mice to cold exposure (4 °C) and monitored core body temperature using implanted telemetry probes. Compared to controls, which showed a transient increase in core body temperature likely reflecting an acute sympathetic thermogenic response, *Orai1* BKO mice exhibited a significantly greater decline in core body temperature (Fig. 3a), indicating impaired thermogenic adaptation. Consistently, infrared thermography revealed lower surface body temperature, particularly in the interscapular BAT region of *Orai1* BKO mice (Fig. 3b). Indirect calorimetry under cold exposure revealed significant reduced oxygen consumption and heat production in *Orai1* BKO mice, with no changes in locomotor activity (Fig. 3c), indicating intrinsic defects in BAT thermogenesis.

**Fig. 3.**
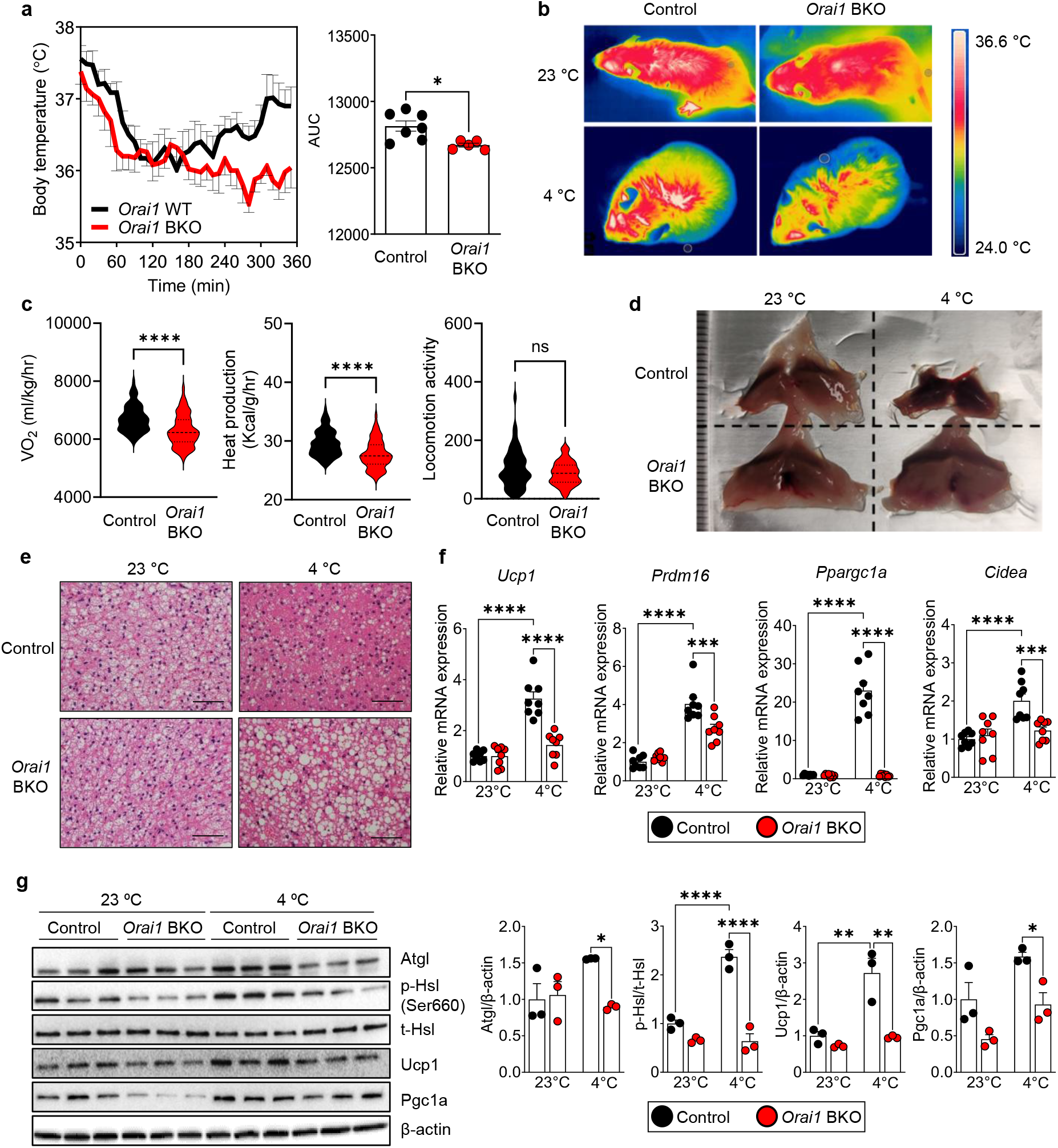
Orai1 deficiency impairs adaptive thermogenesis and lipid mobilization during cold exposure. **a** Core body temperature recording in control and *Orai1* BKO mice subjected to acute cold exposure (4 °C) for 6 hrs. **b** infrared thermography images upon cold exposure. **c** indirect calorimetry during cold exposure (n=6 mice per group). **d** Gross images of BAT of control and *Orai1* BKO under room temperature and cold conditions. **e** H&E staining of BAT. **f** qRT-PCR of thermogenic gene expression in BAT. **g** Western blot analyses of lipolytic and thermogenic proteins. Data are presented as mean ± SEM. Statistical significance: \**p* < 0.05, \*\**p* < 0.01, \*\*\**p* < 0.001, \*\*\*\**p* < 0.0001.

Morphologically, BAT from cold-exposed control mice appeared shrunken and darker, indicative of active lipid mobilization, whereas BAT from *Orai1* BKO mice retained a pale, lipid-rich appearance (Fig. 3d). H&E staining further demonstrated that cold-exposed control BAT contained compact multilocular lipid droplets, while *Orai1* BKO BAT displayed enlarged unilocular ones (Fig. 3E and Supplementary Fig. 3a), consistent with impaired lipid mobilization. No significant histological differences were observed in iWAT under the same conditions (Supplementary Fig. 3b). In line with these morphological changes, expression of thermogenic genes was significantly downregulated in *Orai1* BKO BAT following cold exposure (Fig. 3f). Protein levels of lipolytic markers (Atgl, and phosphorylated Hsl) and thermogenic regulators (Ucp1 and Pgc1a) were lower in *Orai1* BKO BAT (Fig. 3g), indicating impaired activation of lipid mobilization and thermogenic programs. Although thermogenic gene expression remained inducible in *Orai*1 BKO BAT, the magnitude of induction was significantly smaller than in control, consistent with a constraint imposed by reduced basal thermogenic capacity.

To further assess whether Orai1 also mediates thermogenesis in response to adrenergic signaling independently of cold stress, we administered the β_3_-adrenergic agonist CL-316,243 at room temperature. While control mice exhibited a robust increase in core body temperature, *Orai1* BKO mice displayed a significantly blunted thermogenic response (Supplementary Fig. 3c). Moreover, β_3_-adrenergic stimulation failed to induce *Ucp1* and *Ppargc1a* expression in *Orai1* BKO (Supplementary Fig. 3d), indicating that Orai1 contributes to adrenergic-driven thermogenic activation. These data indicate that Orai1 deficiency impairs thermogenic capacity in BAT under both cold and adrenergic stimulation.

### Orai1-dependent Ca^**2+**^ **influx enables cAMP-PKA signaling and mitochondrial uncoupling**

We next investigated the involvement of the cAMP-PKA pathway to elucidate how Orai1 regulates thermogenesis. Cold exposure increased phosphorylation of PKA subunit and downstream substrates in control BAT, whereas this response was markedly blunted in *Orai1* BKO BAT mice (Fig. 4a), indicating that Orai1 contributes to cAMP-PKA activation during thermogenic stimulation. Consistently, intracellular cAMP concentrations were significantly reduced in BAT of cold-exposed *Orai1* BKO mice (Fig. 4b). This reduction was accompanied by the loss of cold-induced transcriptional upregulation of *Adcy3*, a Ca^2+^-stimulated isoform of AC^26^ (Fig. 4c). Expression of other ACs and phosphodiesterases was either unaffected or variably dysregulated in *Orai1* BKO mice (Supplementary Fig. 4a), indicating a selective impairment in cAMP amplification.

**Fig. 4.**
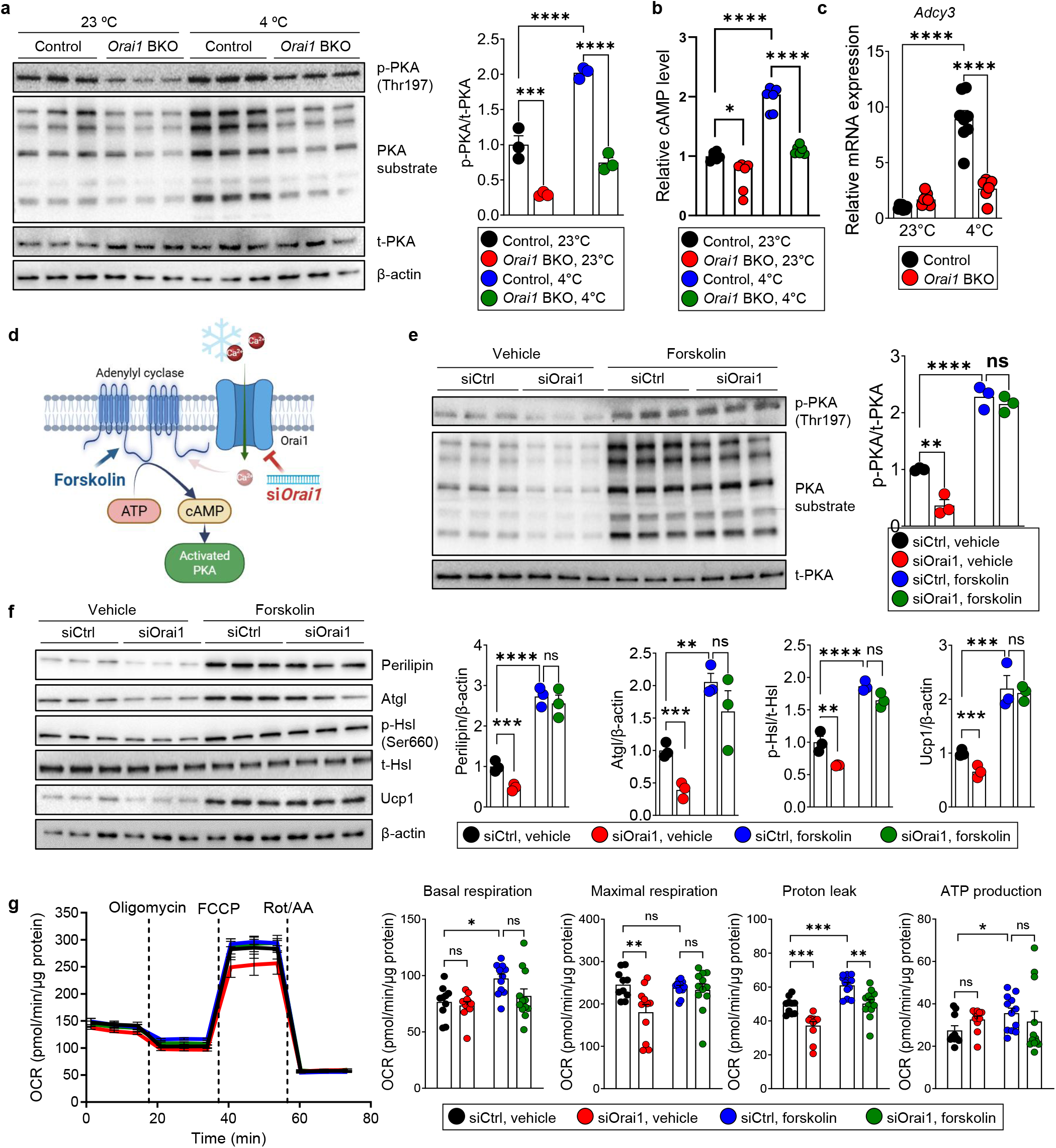
Orai1-dependent Ca^2+^ influx supports full activation of cAMP-PKA signaling and mitochondrial uncoupling. **a** Western blot analysis of PKA and its substrates in BAT from control and *Orai1* BKO under room temperature and cold exposure (4 °C) for 6 hrs. **b** Measurement of intracellular cAMP levels in BAT. **c** *Adcy3* mRNA expression levels in BAT. **d** Experimental design for forskolin rescue assay in *Orai1*-knockdown brown adipocytes. **e** Western blot analyses of PKA activation by forskolin. **f** Western blot analyses of lipolytic proteins and Ucp1 by vehicle- and forskolin-treated cells. **g** Oxygen consumption rate (OCR) measurements in *Orai1*-knockdown cells with forskolin treatment. Data are presented as mean ± SEM. Statistical significance: \**p* < 0.05, \*\**p* < 0.01, \*\*\**p* < 0.001, \*\*\*\**p* < 0.0001.

To further dissect the signaling hierarchy, we performed *in vitro* experiments using primary brown adipocytes with *Orai1* knockdown (Fig. 4d and Supplementary Fig. 4b, c). The expression of *Orai2* and *Orai3* was not altered by *Orai1* knockdown (Supplementary Fig. 4b). In the absence of exogenous stimulation, *Orai1* knockdown reduced PKA activation and phosphorylation, mirroring the *in vivo* phenotype. Notably, treatment of forskolin, a direct AC activator, rescued PKA activity in both control and *Orai1*-knockdown cells (Fig. 4e), consistent with a model in which Orai1 functions upstream of cAMP generation. At the protein level, cold-stimulated expression of lipolytic enzymes and the thermogenic marker Ucp1 was reduced in *Orai1*-knockdown cells, but again, these defects were rescued by forskolin (Fig. 4f). Importantly, genetic rescue by re-expression of ORAI1 in siOrai1-treated human brown adipocytes also restored PKA signaling and HSL phosphorylation (Supplementary Fig. 5a, b), supporting that a role for Orai1 in regulating cAMP-PKA axis.

Mitochondrial function was evaluated by extracellular flux analysis to assess how Orai1 affects respiration and uncoupling capacity. In control brown adipocytes, forskolin enhanced basal respiration, maximal respiration, and proton leak. While basal and maximal respiration were restored in *Orai1*-knockdown cells by forskolin, proton leak remained impaired (Fig. 4g), suggesting that Orai1 also contributes to mitochondrial uncoupling through a Ca^2+^-dependent, cAMP-independent mechanism. Supporting these findings, brown adipocytes from *Orai1* BKO showed impaired PKA phosphorylation and Ucp1 expression even after forskolin stimulation (Supplementary Fig. 4d), and norepinephrine-induced thermogenic gene expression was also blunted (Supplementary Fig. 4e). These results suggest that Orai1 contributes to both cAMP-PKA signaling and mitochondrial activation pathways for thermogenic functions.

### Orai1-mediated Ca^2+^ influx is assocated with mitochondrial Ca^2+^ uptake, remodeling, and thermogenic activation in brown adipocytes

To determine whether Orai1-mediated Ca^2+^ influx directly regulates mitochondrial thermogenic activation (Fig. 5a), we investigated mitochondrial Ca^2+^ dynamics in primary brown adipocytes. Live-cell imaging using Rhod-2 revealed a marked reduction in mitochondrial Ca^2+^ uptake in *Orai1*-silenced brown adipocytes under both SOCE- and cold-stimulated [Ca^2+^]_i_ rise (Fig. 5b, c), indicating that Ca^2+^ influx via Orai1 contributes to mitochondrial Ca^2+^ accumulation. To assess the translational relevance, we examined cold-induced Ca^2+^ response in human brown adipocytes. Acute cold exposure increased cytosolic Ca^2+^ levels in control cells, whereas siRNA-mediated *ORAI1* knockdown significantly attenuated this response (Supplementary Fig. 6a). Importantly, cold-induced mitochondrial Ca^2+^ uptake was also markedly reduced upon *ORAI1* silencing (Supplementary Fig. 6b), indicating that coupling between plasma membrane Ca^2+^ entry and mitochondrial Ca^2+^ signaling is conserved in human brown adipocytes. Mitochondrial structure was examined by transmission electron microscopy, which revealed that cold-exposed *Orai1* BKO BAT exhibited reduced cristae density and disorganized mitochondrial morphology following cold exposure (Fig. 5d and Supplementary Fig. 6c). These ultrastructural abnormalities suggest impaired mitochondrial remodeling during thermogenic adaptation. At the molecular level, expression of mitochondrial oxidative phosphorylation (OXPHOS) complex proteins was significantly reduced in *Orai1* BKO BAT following cold exposure (Fig. 5e). Similarly, *Orai1* knockdown in preadipocytes impaired the induction of OXPHOS components during brown adipocyte differentiation (Fig. 5f), along with reduced expression of Ucp1 and Ppargc1a (Supplementary Fig. 6d). These findings indicate that Orai1 contributes to mitochondrial remodeling and thermogenic gene expression during brown adipocyte maturation and activation.

**Fig. 5.**
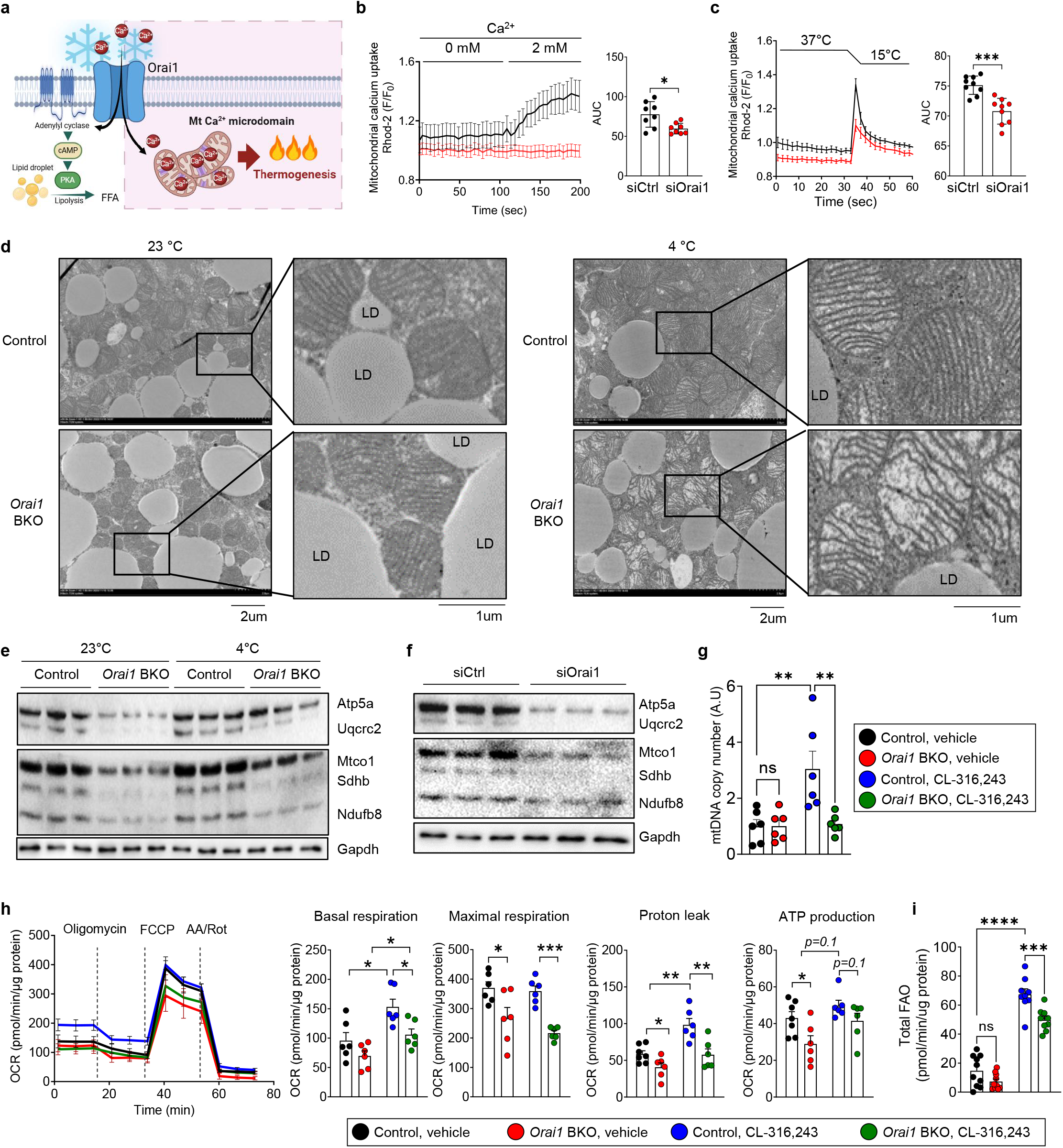
Orai1 deficiency impairs mitochondrial Ca^2+^ uptake and structural remodeling during thermogenic activation. **a** Model illustrating that cold-induced Ca^2+^ influx via Orai1 is essential for mitochondrial Ca^2+^ uptake and subsequent thermogenesis. **b** Rhod-2 fluorescence imaging of mitochondrial Ca^2+^ in siCtrl and siOrai1 cells. **c** Quantification of mitochondrial Ca^2+^ uptake following temperature shifts (37 °C to 15 °C). **d** Transmission electron microscopy images of BAT from 6 hr cold-exposed control and *Orai1* BKO mice. **e** Western blot analysis of OXPHOS proteins in control and *Orai1* BKO BAT after cold exposure (4 °C) for 6 hrs. **f** Western blot analysis of OXPHOS proteins during differentiation of preadipocytes to brown adipocytes treated with siOrai1. **g** Mitochondrial DNA copy number measured in control and *Orai1*-deficient cells with CL-316,243 treatment. **h** OCR analysis of *Orai1*-deficient adipocytes treated with CL-316,243. **i** FAO assay of *Orai1*-deficient adipocytes treated with CL-316,243. Data are presented as mean ± SEM. Statistical significance: \**p* < 0.05, \*\**p* < 0.01, \*\*\**p* < 0.001, \*\*\*\**p* < 0.0001.

Mitochondrial content and functional capacity were further evaluated by assessing mitochondrial DNA copy number in primary brown adipocytes after β_3_-adrenergic stimulation. While CL-316,243 treatment significantly increased mitochondrial content in control cells, *Orai1* BKO cells failed to show such induction (Fig. 5g). Extracellular flux analysis revealed that CL-316,243 treatment enhanced proton leak in control adipocytes. In contrast, *Orai1*-deficient cells failed to increase proton leak, indicating defective thermogenic uncoupling (Fig. 5h). Lastly, to evaluate lipid oxidation capacity, we performed extracellular flux analysis in the presence of etomoxir, an inhibitor of fatty acid oxidation (FAO). While control adipocytes showed a marked increase in FAO upon CL-316,243 treatment, *Orai1* BKO cells exhibited significantly blunted FAO responses (Fig. 5i), further demonstrating impaired mitochondrial fuel utilization. Taken together, these findings indicate that Orai1-dependent Ca^2+^ entry contributes to mitochondrial Ca^2+^ uptake, structural remodeling, uncoupling, and fatty acid oxidation, thereby supporting thermogenic competence in brown adipocytes.

## DISCUSSION

Our study identifies Orai1 as a regulator of Ca^2+^ signaling that orchestrates thermogenic activation in BAT.During thermogenic activation, Orai1 mediates localized Ca^2+^ influx in brown adipocytes, simultaneously activating the cAMP-PKA-dependent lipolytic pathway and promoting mitochondrial Ca^2+^ uptake. These dual actions position Orai1 as a central hub linking Ca^2+^ influx to intracellular thermogenic responses. Importantly, *Orai1* BKO mice already exhibit reduced basal thermogenic capacity under standard housing condition, indicating that Orai1 contributes to basal metabolic homeostasis in brown adipose tissue. Therefore, the blunted response to cold exposure should not be interpreted solely as a defect in cold sensing, but rather as a consequence of diminished thermogenic competence that limits the extent of further activation under cold stress.

Traditionally, SOCE through STIM1/Orai1 interactions has been viewed as the canonical mechanism for Ca^2+^ influx in non-excitable cells^27^. However, our data indicate that Ca^2+^ influx via Orai1 during thermogenic activation does not strictly rely on ER Ca^2+^ store depletion and is only partially attenuated by *Stim1* knockdown, indicating a non-canonical mode of Orai1 activation, with limited STIM1 engagement. Consistently, acute cold increased STIM1 and Orai1 puncta without enhancing oligomerization, further distinguishing this response from classical SOCE. This is consistent with a recent study in sensory neurons showing where cold alone can induce STIM1/Orai1 clustering and activation without ER store depletion or TRP channel involvement^16^ and suggest that cold-induced biophysical alterations in the plasma membrane, including changes in fluidity^28^, lipid raft composition^29^, or rearrangement of cytoskeletal structures, favor Orai1 opening without requiring robust STIM1-Orai1 coupling. A potential limitation of our in vitro system is the use of 15°C as an acute cold stimulus. In vivo, brown adipocytes are unlikely to experience such low temperatures due to tissue insulation and active thermogenesis. Therefore, the temperature shift applied here should be interpreted as an experimental perturbation to induce rapid Ca^2+^ influx rather than a direct physiological mimic of intracellular temperature. Despite this limitation, the consistent Orai1-dependent Ca^2+^ responses observed across in vitro and in vivo models support the functional relevance of this pathway in thermogenic activation. Importantly, we do not interpret these results as evidence for STIM1 dispensability; rather, they indicate that STIM1 engagement during acute cold stimulation is limited compared with canonical SOCE. Notably, our findings distinguish Orai1 from other Ca^2+^ channels like Trpc, Trpm8, and Ryr2, which have been previously implicated in thermosensation or adipocyte Ca^2+^ signaling but appear to play limited roles in cold-induced BAT activation^30-32^. In particular, *Trpm8* knockdown did not diminish cold-evoked Ca^2+^ entry. In contrast, TRPV2 has been directly linked to adaptive thermogenesis rather than cold detection: *Trpv2* knockout mice exhibit cold intolerance, reduced Ucp1 induction, and increased adiposity under HFD^24^.

Mechanistically, we show that Orai1-mediated Ca^2+^ influx contributes to the transcriptional upregulation of *Adcy3*, a Ca^2+^-sensitive isoform of AC enriched in thermogenic adipocytes^33^. This suggests that cytosolic Ca^2+^ may regulate *Adcy3* expression through Ca^2+^/calmodulin-dependent pathway or via Ca^2+^-responsive transcription factors such as CREB or NFAT^34,35^. In turn, increased *Adcy3* expression is consistent with enhances cAMP production and PKA signaling, supporting a model in which Ca^2+^ signaling contributes to amplification of thermogenic pathways. Furthermore, Ca^2+^ entry through Orai1 directly contributes to mitochondrial Ca^2+^ uptake and structural remodeling, thereby supporting cristae integrity, OXPHOS protein expression, and fatty acid oxidation. Thus, Orai1 appears to integrate upstream Ca^2+^ signaling with both the lipolytic and mitochondrial arms of thermogenesis through parallel, Ca^2+^-dependent pathways.

The spatial coupling of Orai1-mediated Ca^2+^ entry with mitochondrial function raises the possibility of localized Ca^2+^ microdomains or membrane-mitochondria contact sites^36^ that enable efficient Ca^2+^ transfer to the mitochondrial Ca^2+^ uniporter (MCU). This model is supported by our findings that mitochondrial Ca^2+^ uptake, OXPHOS expression, and proton leak are impaired in *Orai1*-deficient adipocytes, even in the presence of exogenous cAMP stimulation. The partial rescue of thermogenic gene expression by forskolin, but not mitochondrial uncoupling, suggests an important role of Ca^2+^ influx itself – beyond cAMP elevation – in supporting the full thermogenic program. Our previous work demonstrated that MCU enhances mitochondrial reactive oxygen species production and AMPK activation, facilitating thermogenic gene expression^37^. Here, we extend those findings by identifying Orai1 as a contributor of cytosolic Ca^2+^ influx associated with mitochondrial Ca^2+^ uptake and remodeling. This reveals a functional axis in which Orai1 and MCU cooperate to regulate the thermogenic activation from plasma membrane to mitochondria.

In addition to its acute functional role, we show that Orai1 is transcriptionally upregulated during thergmoenic adaptation. Although the increase in *Orai1* expression under chronic cold conditions, such as those in GSE70437, was modest, its consistency with our acute cold exposure data suggests that Orai1 is regulated both transiently and persistently during thermogenic remodeling. Importantly, this study is the first to utilize a brown adipocyte-specific *Orai1* knockout model, allowing us to dissect the cell-autonomous role of Orai1 in BAT, which had not been captured in previous global or muscle-specific studies^38,39^.

Although *Ucp1* and *Ppargc1a* mRNA levels were relatively preserved under basal conditions, their protein abundance was reduced in *Orai1*-deficient BAT. This discrepancy cannot be fully explained by the present data, but suggests that Orai1 deficiency may influence thermogenic competence through protein-level regulation or post-transcriptional mechanisms in addition to transcriptional regulation. A potential limitation of our model is that *Orai1* deletion occurs during brown adipocyte differentiation via *Ucp1*-*Cre*, raising the possibility that developmental defects may contribute to the observed phenotype. Indeed, *Orai1*-deficient brown adipocytes show reduced mitochondrial content and thermogenic gene expression even at baseline. Nevertheless, post-differentiation perturbations in primary brown adipocytes – siRNA-mediated *Orai1* knockdown and lentiviral ORAI1 re-expression rescue – support a role for ORAI1 in mature cells, partially dissociating the phenotype from developmental history. Therefore, future studies employing temporally inducible Cre models will be necessary to distinguish developmental roles from acute functional requirements of Orai1 in mature BAT. Another limitation of this study is the lack of complete separation between basal thermogenic defects and cold-induced activation, which would be more rigorously addressed by thermoneutral housing experiments. However, the consistent reduction in energy expenditure, thermogenic gene expression, and mitochondrial function observed under baseline conditions strongly supports a model in which Orai1 is required for maintaining thermogenic competence, which in turn constrains adaptive responses to cold.

Importantly, the Orai1-dependent Ca^2+^ signaling identified in mouse brown adipocytes is also recapitulated in human brown adipocytes. From a translational perspective, human BAT is inversely associated with adiposity and insulin resistance, and pharmacological activation of BAT is emerging as a potential strategy to combat obesity and metabolic disease. While no current therapies directly target Orai1, alterations in SOCE components have been observed in obesity and insulin resistance^40^. Our findings suggest that enhancing Orai1 activity in brown adipocytes may offer a novel avenue for promoting thermogenesis, increasing energy expenditure, and alleviating metabolic dysfunction.

In summary, our work reveals that Orai1 functions as a mediator of Ca^2+^-dependent thermogenic signaling in BAT. By coordinating both cAMP-PKA-dependent lipolysis and Ca^2+^-driven mitochondrial activation, Orai1 contributes to thermogenic activation and represents a potential therapeutic target for metabolic diseases.

## ACKNOWLEDGEMENTS

This study was supported by the National Research Foundation of Korea (NRF) grant funded by the Korea government (MSIT) (2023R1A2C1004726, RS-2022-NR070370, and RS-2024-00409403).

## AUTHOR CONTRIBUTIONS

**S.K**.: Conceptualization, Investigation Methodology, Validation, Visualization, Writing – original draft, Writing – review and editing. **N.P.A**.: Investigation, Methodology, Validation, Visualization. **K.P**.: Funding acquisition, Resources, Supervision. **S.C**.: Conceptualization, Funding acquisition, Resources, Supervision, Writing – original draft, Writing – review and editing. **J.N**.: Conceptualization, Funding acquisition, Supervision, Writing – original draft, Writing – review and editing.

## Supplementary Figures

**Supplementary Fig. 1.**
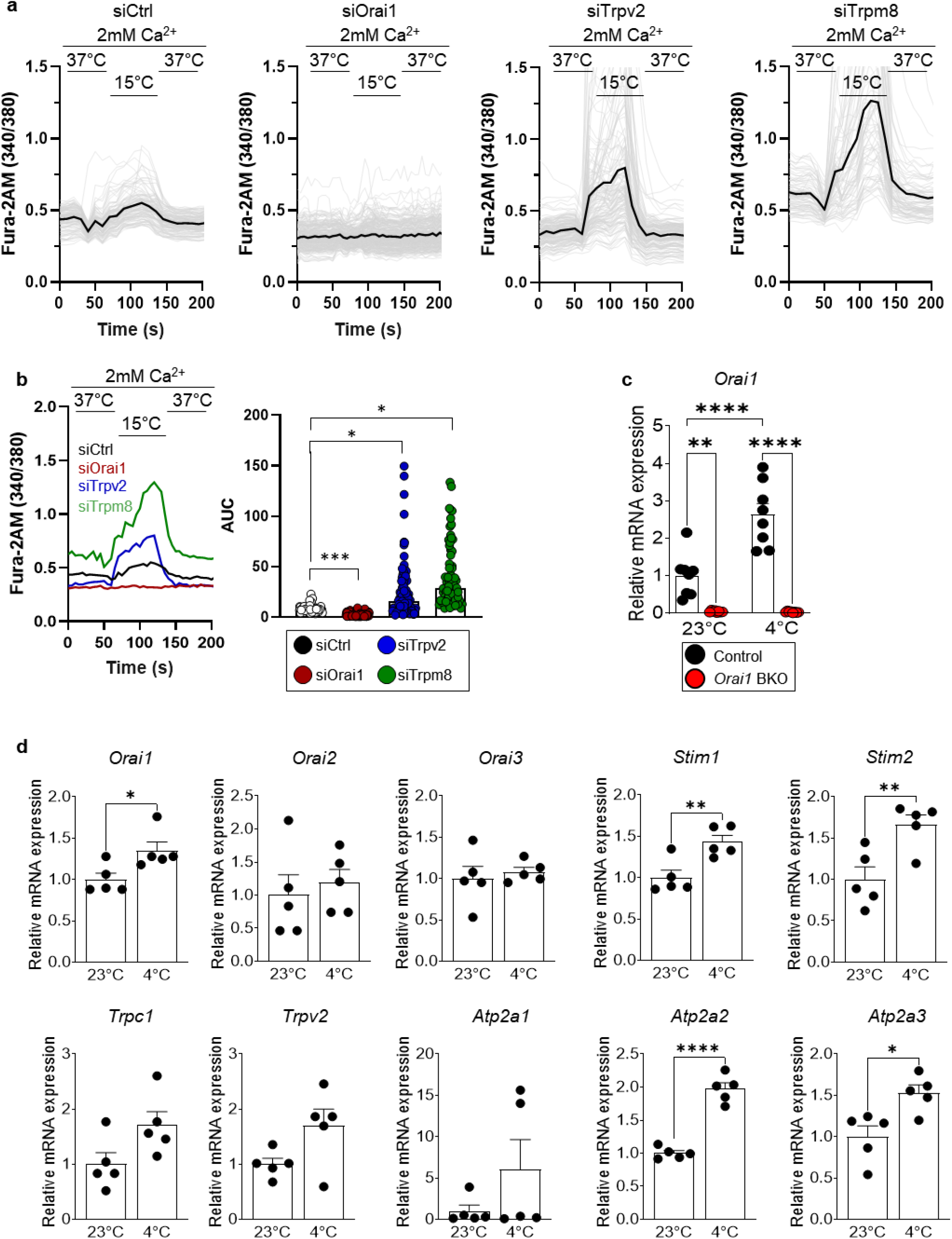
Orai1-related Ca^2+^ entry and its expression in BAT. **a** Intracellular Ca^2+^ levels in primary brown adipocytes with *Orai1, Trpv2*, and *Trpm8* knockdown upon temperature shift from 37 °C to 15 °C. Black lines are average of each traces. **b** AUC analysis of **a. c** Quantification of *Orai1* mRNA expression in brown adipose tissue of *Orai1* control and BKO mice exposed to acute cold exposure (4 °C for 6 hrs). **d** Expressional profiles of SOCE components under chronic cold exposure (3 days), obtained from the GSE70437 dataset. Data are presented as mean ± SEM. Statistical significance: \**p* < 0.05, \*\**p* < 0.01, \*\*\*\**p* < 0.0001.

**Supplementary Fig. 2.**
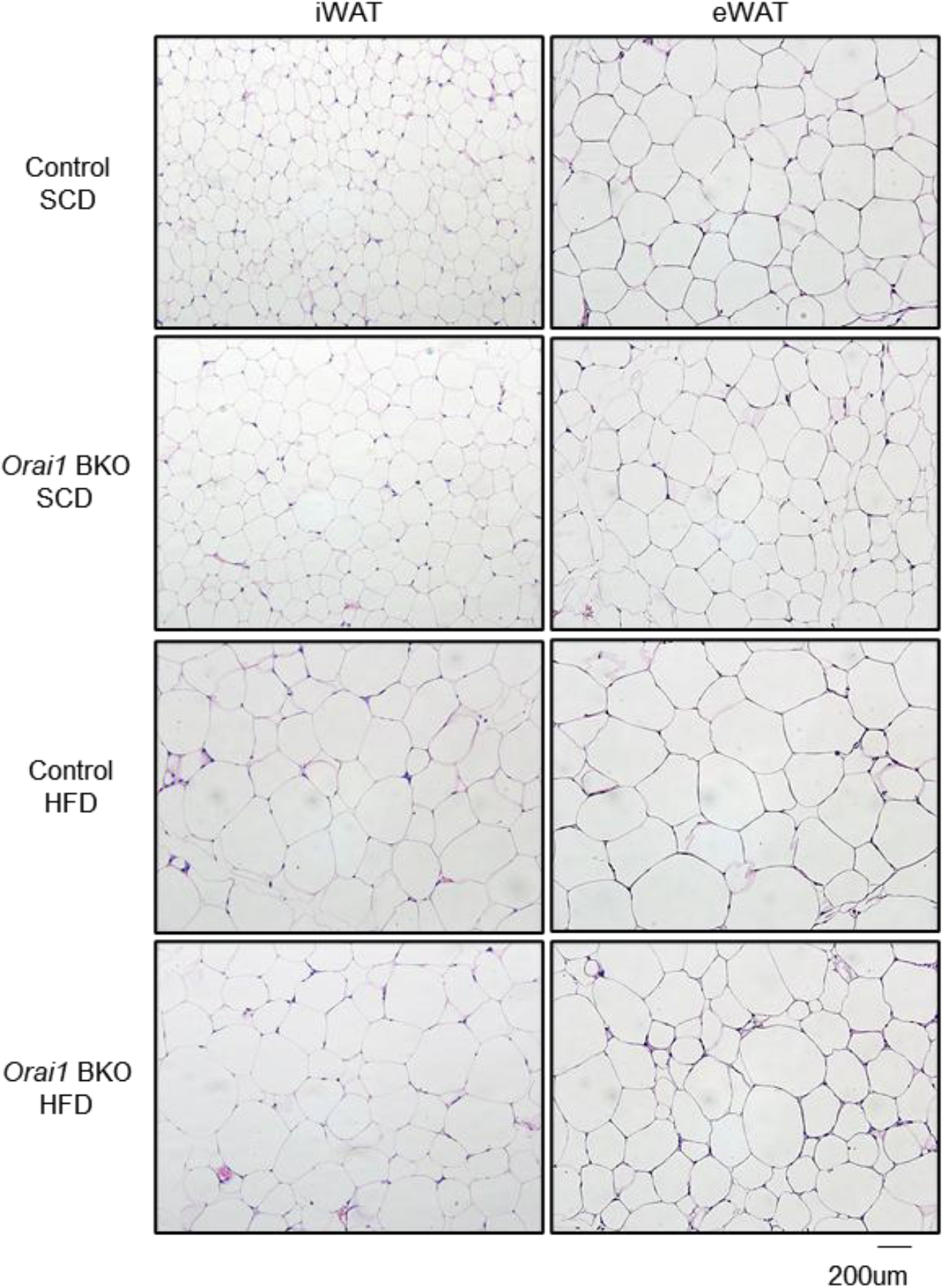
H&E staining of inguinal white adipose tissue (iWAT) and epididymal WAT (eWAT) from control and *Orai1* BKO mice fed with standard chow diet (SCD) or high fat diet (HFD)

**Supplementary Fig. 3.**
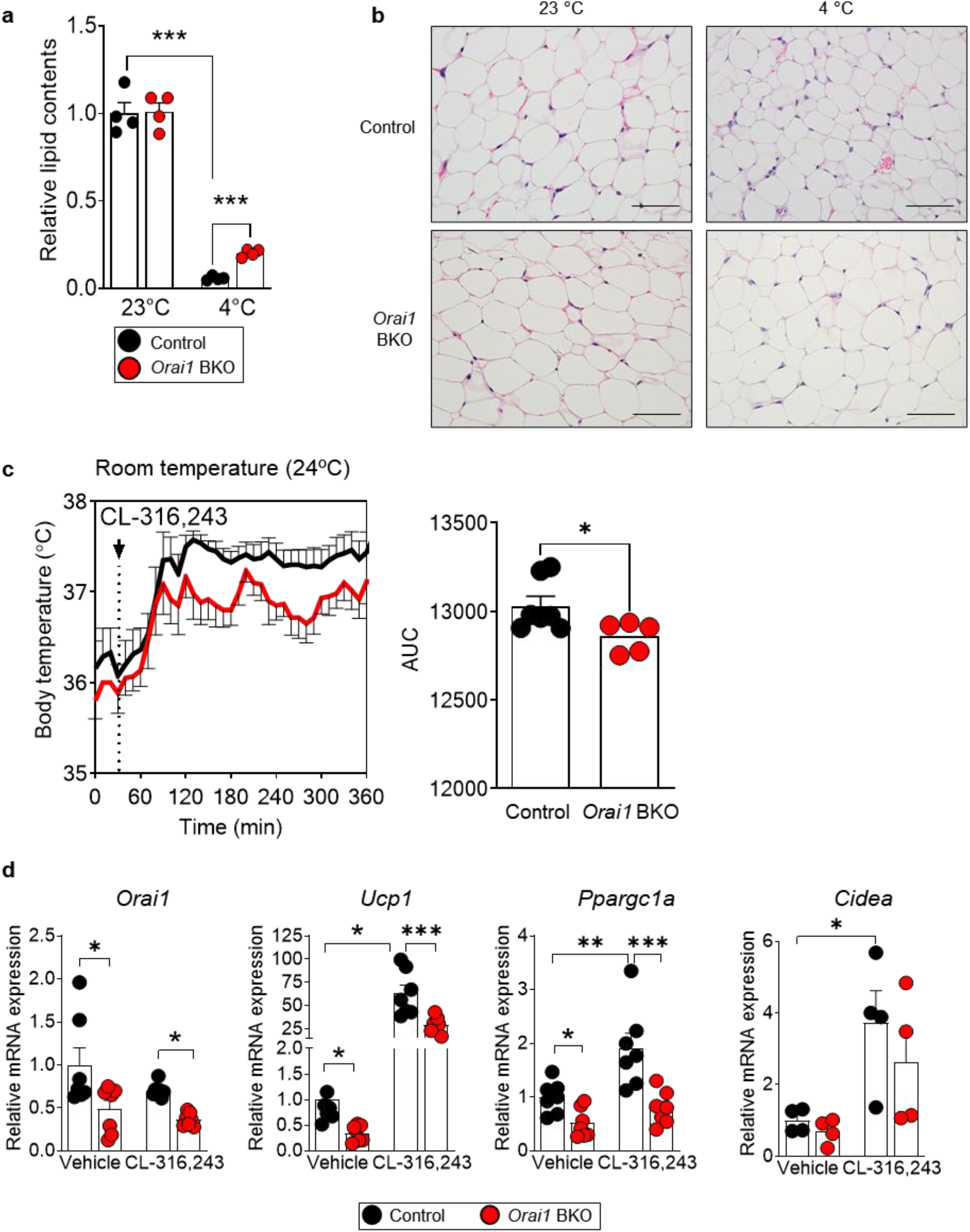
Lipid mobilization and thermogenic responses under cold and β-adrenergic stimulation. **a** Lipid contents of control and *Orai1* BKO BAT from H&E-stained sections (related to Figure 3E). **b** iWAT histology after 4 °C cold exposure (6 hrs). **c** Core temperature measured after CL-316,243 treatment (1 mg/kg i.p.) at room temperature. **d** mRNA expression of thermogenesis genes in primary brown adipocytes following CL-316,243 stimulation. Data are presented as mean ± SEM. Statistical significance: \**p* < 0.05, \*\**p* < 0.01, \*\*\**p* < 0.001.

**Supplementary Fig. 4.**
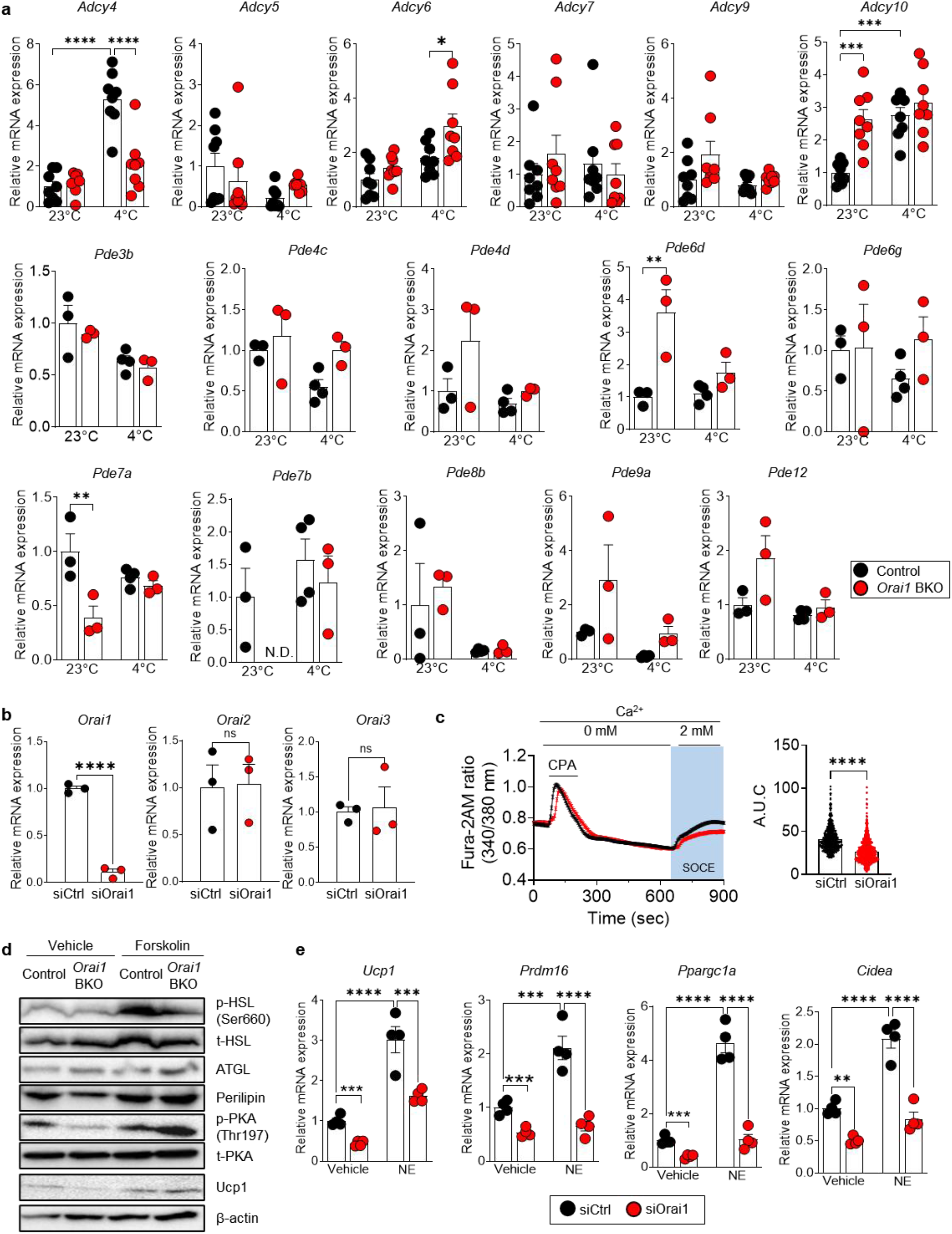
cAMP-related gene expression and signaling in *Orai1*-deficient brown adipocytes. **a** mRNA levels of adenylyl cyclases and phosphodiesterases in BAT from control and *Orai1* BKO with cold exposure (6 hrs). **b** mRNA expression of *Orai1, Orai2*, and *Orai3* in brown adipocytes transfected with siCtrl or siOrai1. **c** Intracellular calcium levels measured using Fura-2 AM in brown adipocytes upon *Orai1* knockdown. CPA (20 μM) was applied as indicated. **d** Western blot of lipolysis and thermogenesis proteins in primary brown adipocytes from control and *Orai1* BKO with forskolin treatment. **e** mRNA expression of thermogenic genes from *Orai1* knockdowned brown adipocytes with norepinephrine (NE) treatment. Data are presented as mean ± SEM. Statistical significance: \**p* < 0.05, \*\**p* < 0.01, \*\*\**p* < 0.001, \*\*\*\**p* < 0.0001.

**Supplementary Fig. 5.**
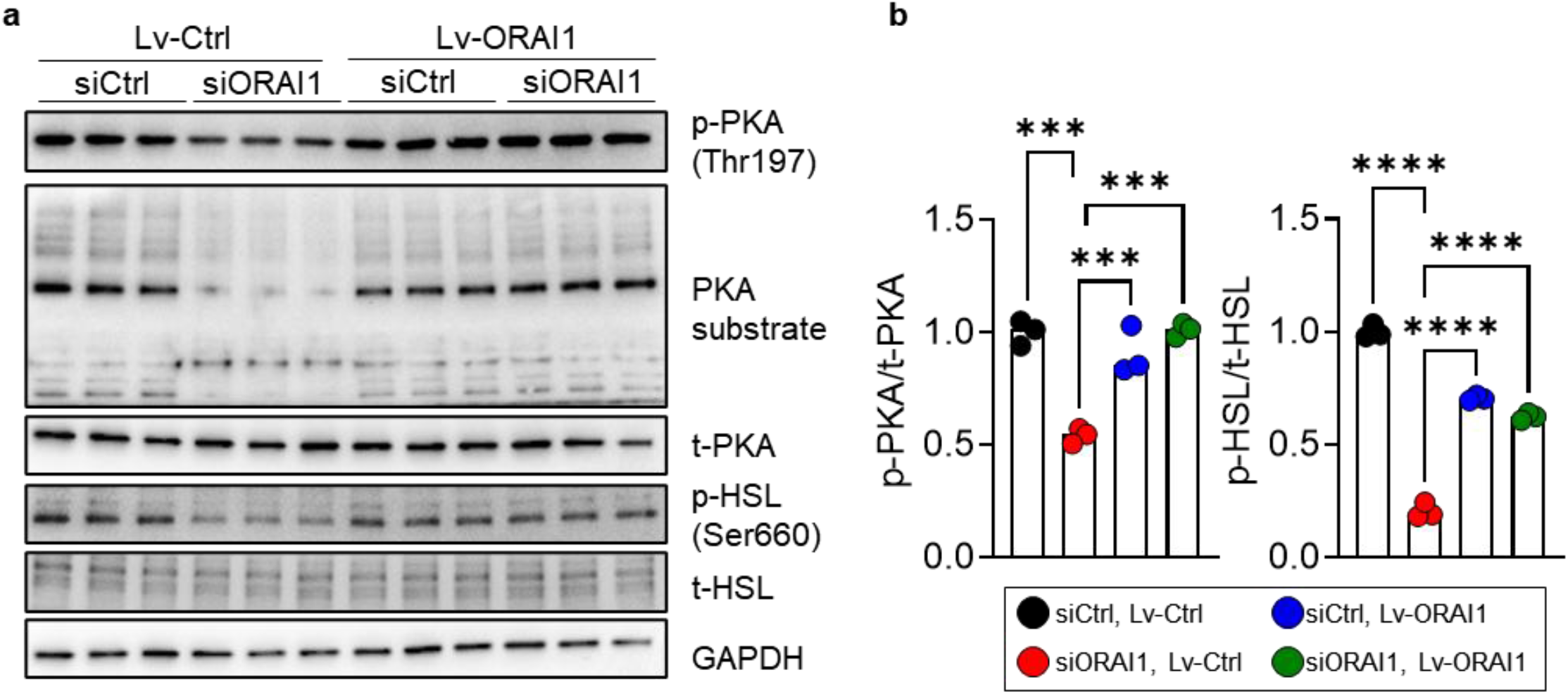
PKA signaling in human brown adipocytes following *ORAI1* knockdown and rescue. **a** Western blot of PKA signaling and HSL phosphorylation under *ORAI1* knockdown followed by lentivirus-mediated *ORAI1* expression in human brown adipocytes. **b** Densitometric quantification from (**a**). Data are presented as mean ± SEM. Statistical significance: \*\*\**p* < 0.001, \*\*\*\**p* < 0.0001.

**Supplementary Fig. 6.**
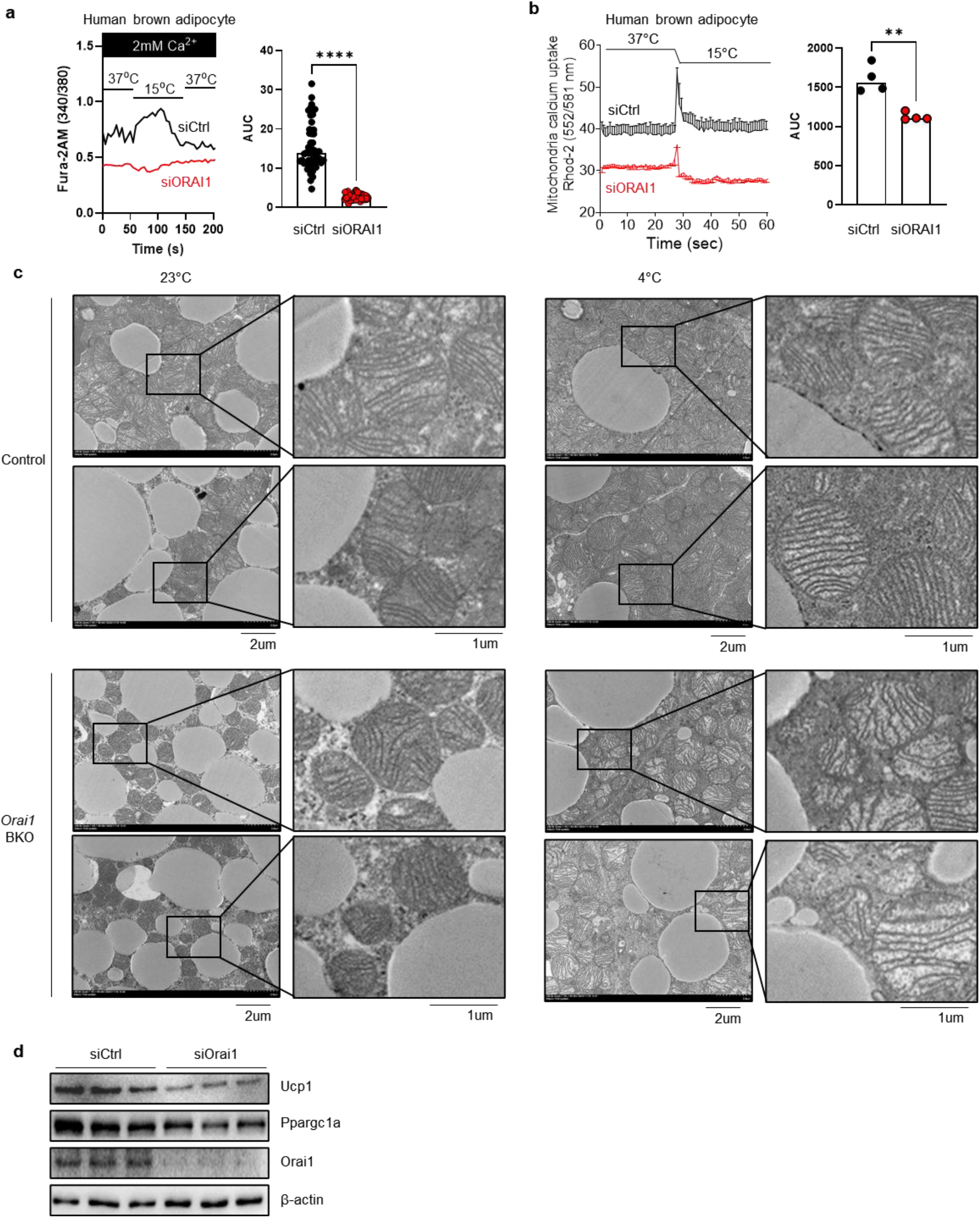
Mitochondrial Ca^2+^ dynamics and ultrastructure in *Orai1*-deficient brown adipocytes. **a, b** Intracellular Ca^2+^ levels (**a**) and mitochondrial Ca^2+^ levels (**b**) in human primary brown adipocytes following temperature shift from 37 °C to 15 °C. **c** Transmission electron microscopy of BAT from control and *Orai1* BKO mice after 6 h cold exposure at 4 °C. Insets show magnified mitochondria. **d** Western blot of thermogenic proteins from brown adipocytes with *Orai1* knockdown.

## Supplementary Tables

**Supplementary Table 1.** List of antibodies used in this study.

| Target protein | Source | Identifier |
| --- | --- | --- |
| Orai1 | Mybiosource | MBS3209798 |
| Ucp1 | Abcam | ab209483 |
| Atgl | Cell Signaling | 2138S |
| HSL (total) | Cell Signaling | 4107S |
| HSL (p-Ser660) | Cell Signaling | 45804S |
| Perilipin | Abcam | Ab3526 |
| Pgc1a | Abcam | ab191838 |
| PKA (total) | Cell Signaling | 4782 |
| PKA (p-Thr197) | Cell Signaling | 4781 |
| PKA substrates | Cell Signaling | 9624S |
| OXPHOS Complex cocktail | Abcam | ab110413 |
| b-Actin | SantaCruz | Sc47718 |
| GAPDH | SantaCruz | Sc47724 |

**Supplementary Table 2.**
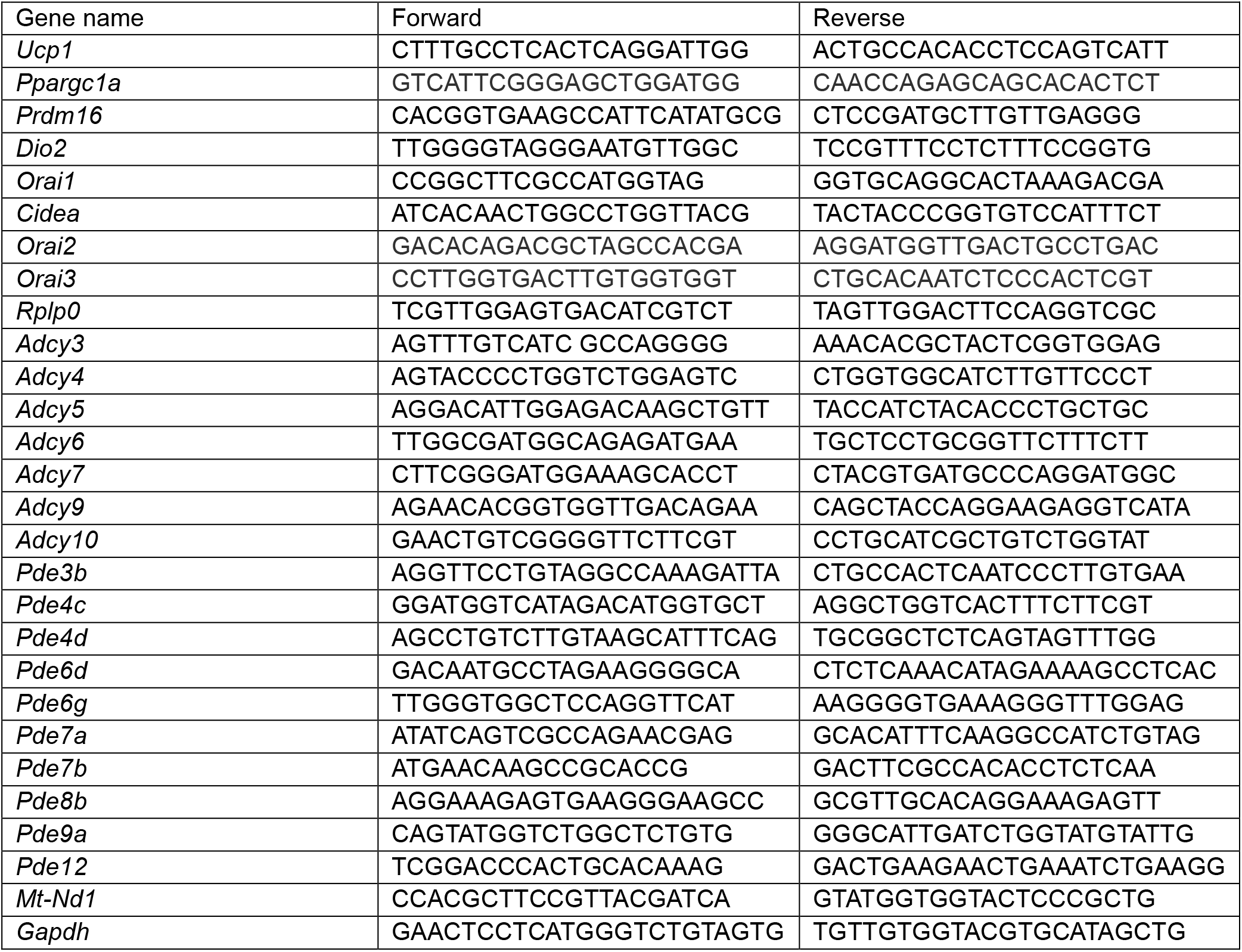
Primer sequences used in this study.

